# High-resolution genomic comparisons within *Salmonella enterica* serotypes derived from beef feedlot cattle: parsing the roles of cattle source, pen, animal, sample type and production period

**DOI:** 10.1101/2020.10.26.356493

**Authors:** Gizem Levent, Ashlynn Schlochtermeier, Samuel E. Ives, Keri N. Norman, Sara D. Lawhon, Guy H. Loneragan, Robin C. Anderson, Javier Vinasco, Henk C. den Bakker, H. Morgan Scott

## Abstract

*Salmonella enterica* is a major foodborne pathogen, and contaminated beef products have been identified as the primary source of *Salmonella*-related outbreaks. Pathogenicity and antibiotic resistance of *Salmonella* are highly serotype- and subpopulation-specific, which makes it essential to understand high-resolution *Salmonella* population dynamics in cattle. Time of year, source of cattle, pen, and sample type(i.e., feces, hide or lymph nodes) have previously been identified as important factors influencing the serotype distribution of *Salmonella* (e.g., Anatum, Lubbock, Cerro, Montevideo, Kentucky, Newport, and Norwich) that were isolated from a longitudinal sampling design in a research feedlot. In this study, we performed high-resolution genomic comparisons of *Salmonella* isolates within each serotype using both single-nucleotide polymorphism (SNP)-based maximum likelihood phylogeny and hierarchical clustering of core-genome multi-locus sequence typing. The importance of the aforementioned features on clonal *Salmonella* expansion was further explored using a supervised machine learning algorithm. In addition, we identified and compared the resistance genes, plasmids, and pathogenicity island profiles of the isolates within each sub-population. Our findings indicate that clonal expansion of *Salmonella* strains in cattle was mainly influenced by the randomization of block and pen, as well as the origin/source of the cattle; that is, regardless of sampling time and sample type (i.e., feces, lymph node or hide). Further research is needed concerning the role of the feedlot pen environment prior to cattle placement to better understand carry-over contributions of existing strains of *Salmonella* and their bacteriophages.

**Importance:** *Salmonella* serotypes isolated from outbreaks in humans can also be found in beef cattle and feedlots. Virulence factors and antibiotic resistance are among the primary defense mechanisms of *Salmonella*, and are often associated with clonal expansion. This makes understanding the subpopulation dynamics of *Salmonella* in cattle critical for effective mitigation. There remains a gap in the literature concerning subpopulation dynamics within *Salmonella* serotypes in feedlot cattle from the beginning of feeding up until slaughter. Here, we explore *Salmonella* population dynamics within each serotype using core genome phylogeny and hierarchical classifications. We used machine-learning to quantitatively parse the relative importance of both hierarchical and longitudinal clustering among cattle host samples. Our results reveal that *Salmonella* populations in cattle are highly clonal over a 6-month study period, and that clonal dissemination of *Salmonella* in cattle is mainly influenced spatially by experimental block and pen, as well by the geographical origin of the cattle.

## Introduction

Globally, *Salmonella enterica* subsp. *enterica* is among the leading foodborne bacterial pathogens and is responsible for negative effects both on public health and on the economy (1). While there are over 2,500 identified *Salmonella* serotypes, only a minor set (~100) of these serotypes are commonly identified as the source of human *Salmonella* infections and outbreaks (2). Each year, approximately 93.8 million human cases occur worldwide due to *Salmonella*, resulting in 155,000 deaths (3). Human infections caused by pathogenic *Salmonella* strains often result in mild gastroenteritis and do not require antibiotic treatment. At-risk individuals, such as infants, the elderly, or immunocompromised patients, may require the use of antibiotics to counter the pathogen’s invasive and adverse health effects. Over the last decade, high levels of hospitalization and increased mortality rates associated with multidrug-resistant (MDR) *Salmonella* serotypes (4, 5) have, indeed, become a serious public health concern (6, 7).

The majority of *Salmonella* outbreaks are related to the consumption of contaminated food products (3). *Salmonella*-contaminated beef products are a major culprit in foodborne S*almonella* outbreaks (8). *Salmonella* contamination of carcass surfaces can occur during slaughter evisceration and de-hiding processes due to direct contact with cattle feces and aerosolization of fecal material from hides, respectively. Lymph nodes harboring *Salmonella* embedded in fat tissue can also contaminate final beef products (e.g., ground beef) as fat trim (9).

Since the gastrointestinal tract of cattle is known to be a natural niche for a number of *Salmonella* serotypes, those that can cause human infections through the ingestion of contaminated beef products are commonly found not only in cattle but also in their feedlot pen environments (10–15).*Salmonella* serotypes Typhimurium, Newport, Enteritidis, Montevideo, Anatum, Cerro, Kentucky, and Mbandaka are commonly found in cattle, beef products, and in cattle production environments (8, 11, 14–18). Among these, Typhimurium and Newport are the serotypes most commonly derived from human cases of *Salmonella*, followed next by Anatum and Montevideo (16, 19, 20). Cattle-origin *S*. Typhimurium and Newport are also serotypes commonly associated with a phenotypic MDR profile (8, 11, 14–17).

In an effort to minimize public-health risks of *Salmonella* originating from cattle, multiple teams of researchers have previously explored serotype-level dynamics of *Salmonella* in healthy cattle among samples taken from feces, lymph nodes and hides, as well as from feedlot pen environments. These teams have documented not only a shift of the serotype-level population as influenced by geographical, ecological, and seasonal (or temporal) changes, but also an elevated association between specific antibiotic resistance profiles found in specific serotypes (10, 15, 21–25)

Pathogenicity (virulence) and antibiotic resistance are two of the major adaptation, survival, and defense mechanisms of *Salmonella.* These mechanisms are located on bacterial chromosomes or plasmids, and are considered among either the core- or accessory-genomes of *Salmonella* (26–32). The majority of the core genome and almost half of the accessory genes of *Salmonella* are serotype-specific (29, 33). For instance, plasmidal defense mechanisms are mainly responsible for the transmission of virulent and antibiotic-resistant strains among humans, animals, and the environment, and are often serotype-specific (34, 35). The increasing use of whole-genome sequencing and the development of allele-and SNP-based techniques used for high-resolution genomic comparisons of *Salmonella* (36–39) recently revealed that pathogenicity and antibiotic resistance mechanisms in *Salmonella* were associated with genetic diversity of clonal groups belonging to the same sequence type, or else to lineages within the same serotype (30, 33, 40–42).

Several genomic epidemiology studies have investigated potential drivers of clonal expansion of outbreak-related serotypes, such as *S*. Typhimurium, Enteritidis, Newport, Dublin, Kentucky, Montevideo, and Cerro, all of which are highly prevalent in cattle, by using genomic comparisons of the isolates derived from various host species, food sources, environments and geographical locations (29, 30, 34, 36, 40–54). A subset of these studies demonstrated that subpopulation-level genetic variations were strongly influenced by time and varied by geographical location at global- (36, 40, 41, 49, 50), country- (49, 51), and state-levels (29, 30, 44, 52). In addition to geographical location, a group of studies also showed distinct host- and source-specific macroevolutionary patterns that were observed across human-, animal-, food-, and environment-origin *Salmonella* isolates (29, 30, 34, 40, 42–46, 53). The genetic variation observed among the geographical location, host and environment were attributed mainly to the alterations observed in the prophage regions (43, 45, 54), the absence or presence of pathogenicity-related genes (36, 42–44, 46, 52, 54) as well as antibiotic resistance genes (41, 44, 50).

In the literature, there remains a gap in studies of high-resolution population dynamics of *Salmonella* serotypes in cattle from the beginning of the feeding period until slaughter; in case of the latter, this represents the most critical period for mitigating *Salmonella* impacts on public health. Notably, the temporal evolutionary changes observed within serotype and the potential factors that play a role in the microevolutionary patterns of *Salmonella* populations in cattle are yet to be discovered and thoroughly understood (4).

Previous work by our group measured the effects of single-dose metaphylactic antibiotic treatments on *Salmonella* prevalence in beef feedlot cattle feces, hides, and lymph nodes, and reported no treatment effects on the prevalence or phenotypic antibiotic resistance profiles of *Salmonella* (25). However, the serotype-level distribution of *Salmonella* (Anatum, Lubbock, Cerro, Montevideo, Kentucky, Newport, and Norwich) that corresponded to a single sequence type was found to be mostly influenced by the geographical origin of the cattle and the experimental blocks of pens that housed them through the study period.

Our primary objective in the present analysis was to investigate the temporal microevolutionary patterns of *Salmonella* within each serotype to quantify the relative importance of cattle source and experimental allocation via blocks and pens over a study period of several months. We also explored the specific role of sample types (i.e., feces, lymph nodes, hides) by using in each case the core-genome SNP-based maximum-likelihood phylogeny and hierarchical clusters of core-genome multilocus sequence types (HierCC)analyses to explore the multiple levels of resolution of core-genomic relationships among the subpopulations within the *Salmonella* serotypes. We further compared the HierCC-levels and cluster numbers with a publicly available *Salmonella* whole-genome database that includes human clinical disease isolates (EnteroBase Platform http://enterobase.warwick.ac.uk/species/index/senterica) to explore the potential public health risks of the *Salmonella* subpopulations identified in our study (38, 55). Secondarily, using a supervised machine-learning algorithm, we explored the relative importance of each experimental component in contributing towards clonal *Salmonella* expansion within each serotype. Lastly, we investigated the presence and distribution of antibiotic resistance genes, along with plasmids and pathogenicity islands, within the entire *Salmonella* population isolated during the study.

## Materials and Methods

This study was reviewed for animal use and biosafety and approved by the West Texas A&M University/Cooperative Research, Educational and Extension Team Institutional Animal Care and Use Committee (Protocol no. 05-09-15) and the Texas A&M University Institutional Biosafety Committee (IBC2017-049), respectively.

The experimental design of the study was previously described by Levent et al. (2019) (25). Briefly, a total of 134 cross-bred beef cattle was purchased from two different sources (35 cattle from Source 1 [Hereford, Texas], and 99 cattle from Source 2 [Abilene, Texas], located 480 km apart from each other) in west Texas, USA. Cattle were shipped to a research feedlot operated by West Texas A&M University and located near Canyon, Texas. Upon arrival to the feedlot, cattle were source- and weight-blocked and allocated into four blocks (1-4); within each block, cattle were randomized into three pens to control for possible cattle source- and host-related confounders. Later, cattle pens (with 11-12 steers) in each block were randomly assigned to receive: 1) ceftiofur, 2) tulathromycin or, 3) remain as a control group without any antibiotic treatment, respectively. For each treatment pen, all but one (or two) of the cattle were administered subcutaneously a single dose one-time injection of either ceftiofur crystalline-free acid (Excede^®^, Zoetis Inc., Kalamazoo, MI, USA) at 6.6 mg/kg or else tulathromycin (Draxxin^®^, Zoetis Inc., Kalamazoo, MI, USA) at 2.5 mg/kg. Staff involved in this study were blinded as to treatment to avoid possible selection bias. From time of antibiotic treatment (Day 0), until the time of slaughter (between Days 99 - 141), cattle were fed and raised without any additional antibiotic supplements. Starting from Day 99, cattle that had reached the desired body weight for slaughter were sent to the slaughterhouse as experimental blocks until Day 141, which was the end of the study. Fecal samples (*per rectum*) were collected via sterile obstetric gloves immediately prior to the antibiotic treatment (Day 0), a week after the injection (Day 7), and on multiple additional days up until slaughter. At slaughter, one final fecal sample and an additional hide swab, rubbed from a one-m^2^ cranial ventral (brisket) area, were collected from each steer. In addition, two peripheral sub-iliac lymph nodes were collected from each hot carcass during slaughter at a local federally inspected processing plant. Study design details relating to allocation of the 134 cattle purchased from two different locations (sources) to the experimental blocks and pens and the sample types corresponding to each of the sampling days are provided in Table 1.

**Table 1.** Study design representing number of cattle distributed among source, block, and pen along with the sample types collected each sampling day. Cattle in Pens 7, 8 and 9 were from Source 1 and were located next to one another in the feedlot. Cattle from Source 2 were located next to one another in Pens 51, 52, 53, 54, 55, 56, 57, 58, and 59. A schematic of the pen locations in the feedlot was published previously in Levent et al. (2019) (25). Pens within each block (see Table) were randomly assigned for cattle to receive either ceftiofur (Cef) or tulathromycin (Tul) on Day 0 prior to first fecal sample collection, or else to remain as control (Cont) animals.

| Source | Block | Pen identifier | Treatment | Number of cattle per pen | Feeding period (feces only) | Slaughter period (feces, lymph node, hide) |
| --- | --- | --- | --- | --- | --- | --- |
| Hereford, TX (Source -1) | 1 | 7 | Tul | 12 | Day 0 and 7 | Day 134 |
|  |  | 8 | Cont | 11 |  |  |
|  |  | 9 | Cef | 12 |  |  |
| Abilene, TX (Source-2) | 2 | 51 | Tul | 11 |  | Day 141 |
|  |  | 52 | Cont | 11 |  |  |
|  |  | 53 | Cef | 11 |  |  |
|  | 3 | 54 | Cont | 11 |  | Day 120 |
|  |  | 55 | Cef | 11 |  |  |
|  |  | 56 | Tul | 11 |  |  |
|  | 4 | 57 | Cef | 11 |  | Day 99 |
|  |  | 58 | Tul | 11 |  |  |
|  |  | 59 | Cont | 11 |  |  |

*Salmonella* isolates derived from fecal samples during the early feeding period (Day 0 and 7) and fecal, sub-iliac lymph node, and hide swab samples obtained at slaughter included for analysis in this study are previously reported in detail by Levent et al. (2019). A total of 399 isolates was previously isolated, sequenced, serotyped and sequence typed (using legacy 7-gene MLST [multi-locus sequence type]), and were identified as Lubbock ST413 (n =136), Anatum ST64 (n=113), Montevideo ST138 (n=68), Cerro ST138 (n=64), Kentucky ST152 (n=11), Newport ST118 (n=6), and Norwich ST2119 (n=1).

The methods related to *Salmonella* isolation, DNA extraction, whole-genome sequencing, genome quality, genome assembly, and *in silico* tools used for legacy 7-gene MLST and serotyping are previously published by Levent et al., 2019 (25). Paired-end raw sequencing reads and assemblies of 399 *Salmonella* isolates have previously been deposited in the NCBI database under BioProject number PRJNA521731. BioSample numbers of each assembly, and the metadata related to each isolate are listed in Supp. Dataset 1.

### Phylogenetic analyses

*Salmonella* strains restricted to those from serotypes Anatum, Cerro, Montevideo, and Lubbock and isolated from multiple sample types arising from individual cattle throughout the study were included in the core-genome SNP analysis using maximum-likelihood phylogeny. This was performed to obtain a high degree of resolution of the population structure within each serotype; importantly, all isolates of one serotype were identified as being from the same legacy 7-gene MLST group (39). Cattle source, block, pen, day, and sample type distribution of the isolates included in the SNP-based phylogenetic analysis from each of the four serotypes are presented in Table 2. Kentucky, Newport, and Norwich isolates were excluded from phylogenetic analysis due to small sample sizes (n<12).

**Table 2.**
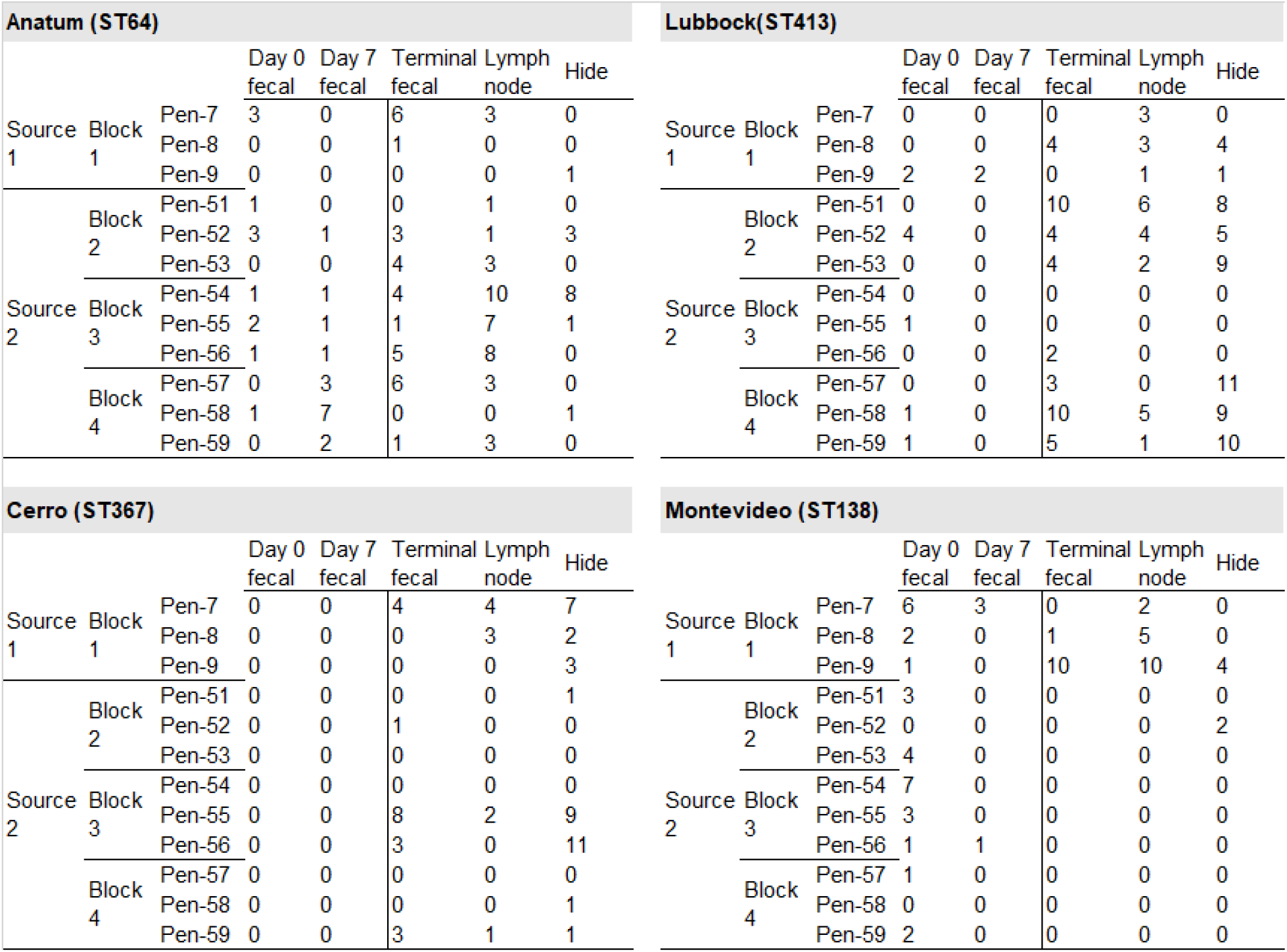
Number of isolates selected for phylogenetic analysis by serotype, source, block, pen, day, and sample type.

### Reference genome selection

In order to perform core-genome-level SNP analysis, the complete reference genomes at the closest genomic distance for each serotype were selected using the Similar Genome Finder Service on PATRIC (the Pathosystems Resource Integration Center: available at https://www.patricbrc.org/app/GenomeDistance). The selection criteria were kept at default threshold values (maximum hit value of 50, *P* value of 1, and Mash/MinHash distance [estimating the distance based on the rate of sequence mutation] value of 0.05) (56, 57). The selected reference genomes were further screened for prophage regions using PHASTER (Phage Search Tool Enhanced Release, available at http://phaster.ca/) (58). The complete and questionable prophage regions were further masked using BEDTools v.2.18 (59) and were carried to the next step; that is, alignment of sequencing reads from the isolates for phylogenetic analyses.

### Variant calling, tree inference and tree visualization

Reference alignment and variant-calling processes for those isolates that shared the same serotype were performed using the McOutbryk SNP calling pipeline (available under the Massachusetts Institute of Technology license at https://github.com/hcdenbakker/McOutbryk) using raw (FASTQ) sequencing short reads with a minimum sequencing depth of 24X and a reference genome (60). This pipeline uses the McCortex tool (61) to build graphs of a reference sequence along with the data of the genomes to be queried for SNP calling. The SNP-calling stage consists of two phases: (Phase 1) an initial phase, which consists of a comparison of the reference graph and each query genome to construct a list of putative variable sites within the population, and (Phase 2) a final SNP-calling phase, which calls the allele for each putative SNP site found in Phase 1. While the SNP calling is done *de novo* at the initial step, the pipeline uses BWA (Burrow-Wheeler aligner)-mem (62) at the final step to place the SNP sites in relation to the reference sequence. In addition to BWA, the pipeline relies on VCFtools (63), and vcflib (available under the MIT license at https://github.com/vcflib/vcflib) for VCF (Variant Call Format) file manipulation. K-mer size 33 was set for variant analysis, and highly divergent isolates (number of SNPs > 5000) were excluded from the analysis by default. The matrices containing SNP sites were later evaluated for the best-fit nucleotide substitution model using IQ-tree v.1.6.10 with ascertainment bias correction (ASC) option to construct a maximum-likelihood phylogeny (64). Later, the phylogenetic trees were inferred using the selected nucleotide submission model, including bootstrap estimates obtained with 1,000 iterations (64). The resulting trees were visualized and annotated with cattle source, pen, block, day and sample type as features of the isolates using the interactive tree of life (iTOL v.4) (65). The number of SNPs corresponding to the tree scale and the strong branch support values (bootstrap values of 80%-100%) of the clades were presented on each resulting phylogenetic tree.

### Hierarchical clustering of cgMLST and feature importance analyses

Hierarchical clustering of the cgMLST (core-genome multilocus sequence typing, also known as HierCC) cluster number of the isolates – including all serotypes – were determined at multiple levels of HierCC clusters using the EnteroBase platform (based on cgMLST V2 + HierCC V1) to obtain high resolution genomic-based comparisons via HierCC (38, 55). The HierCC cluster numbers observed for each of the isolates derived from our study were also compared with the publicly available whole-genomes in the *Salmonella* database provided by EnteroBase to explore the public health-related significance of our findings at the HC0-,HC2-, HC5-, HC10- and HC50-levels (38).

Furthermore, the importance of isolate features such as, cattle source (1 and 2), block (1, 2, 3, and 4), day (0, 7, and 122 [collapsed to the average day from slaughter days 99, 120, 134, and 141]) and sample type (fecal, lymph node, and hide) for the prediction of the indistinguishable isolates (i.e., those that were highly clonal and designated to the same HC0 cluster; that is, with no cgMLST allele difference) were explored for each serotype population using the Orange v.3.26.0 data-mining toolbox (66). The supervised machine learning algorithm Random Forest (67, 68) was used to build a set of (n=10) decision trees for the clonal groups using the features listed above, and the resulting trees were explored using Pythagorean Forest (69) algorithm. The best tree that that requires only a few attributes to split the branches was selected to evaluate the feature importance on the tree. Importance of the features was ranked from the most important to the least important based on the values predicted by the ReliefF algorithm (70, 71).

### Characterization of antibiotic resistance genes, plasmids, and pathogenicity islands

*In silico* plasmid and antibiotic resistance gene detection and identification of 399 assembled genome sequences were performed using ABRicate v.0.8.7 (https://github.com/tseemann/abricate) using the ResFinder (72) and the PlasmidFinder (73) databases as well as the SPIFinder (74) database that were obtained from the Center for Genomic Epidemiology server (available online at https://bitbucket.org/genomicepidemiology/spifinder_db/src/master/).

## Results

### Population structure of *Salmonella*

*Salmonella* Anatum, Cerro, Montevideo, and Lubbock isolates collected from individual steers throughout the study period were included in core-genome SNP-based phylogenetic analyses. Based on the resulting trees, the highest number of core-genome SNPs (n=84) was observed among Lubbock isolates, followed by Anatum (66 SNPs), Cerro (17 SNPs) and Montevideo (12 SNPs); importantly, each demonstrated distinct cattle source-, block-, pen-, and day-related patterns.

Population-based HierCC analysis showed that there were no more than 100 cgMLST allelic distance between *Salmonella* genomes of each serotype. Based on HierCC designations, the highest level of cluster variation across all sample features and within the constraints of a 122-day study was observed in Anatum (HC100), Lubbock (HC10) and then Cerro (HC10) isolates, followed by Montevideo (HC5), Kentucky (HC5), and Newport (HC0). Details concerning the multi-level HierCC cluster numbers for the isolates found in this study are presented in Supp. Dataset 1 and Fig. S1.

As of August 21, 2020, there were 7,149 publicly available *Salmonella* genomes in EnteroBase with an allelic distance equal or less than 50 (HC50) found to be no; of those, 6,934 genomes were found in the same HC50 cluster of serotypes, followed by 145 genomes in HC20, 65 genomes in HC10, and 4 genomes in the same HC5 cluster. A single isolate was found in the HC2 cluster suggesting nearly complete homology. Detailed information related to the HierCC cluster numbers, along with the geographical region of origin, source type and other *Salmonella* genome-related data were downloaded from EnteroBase and provided in Supp. Dataset 2. Technical information related to the HierCC clustering method is also provided in the EnteroBase (available at https://enterobase.readthedocs.io/en/latest/features/clustering.html accessed on 10/20/2020)

Similarly, serotype-specific population structures of *Salmonella* isolates are presented in further detail within specific sections immediately below.

### *S*. Lubbock

The complete genome of a single previously reported Lubbock isolate (GenBank: CP032814.1) was found to be the closest genome match to those Lubbock isolates derived from our study. A total of 11 prophage regions were detected and masked in this reference genome (Table S1). The phylogenetic tree of 136 Lubbock isolates was inferred using the K2P+ASC model, which was the best nucleotide substitution model according to the BIC value. The resulting tree had two clades (Clade I and II) and well-supported subclades (bootstrap values of > 80%) with distinct patterns for source, block, pen, and day (Fig. 1). While the isolates from Block 4 were commonly observed in Clade I, isolates from Block 1 (Source 1) were only observed in Clade II. Lubbock isolates derived from the early feeding period (Days 0 and 7) were only observed in Clade II; that is to say, there was no early feeding period isolates observed in Clade I. The phylogeny of Lubbock isolates yielded the highest total number of SNPs (n = 84) when compared to the total number of SNPs obtained from other serotypes included in this study. There were no sample type-related (feces, lymph node and hide) phylogenetic differences observed among the Lubbock isolates derived at slaughter.

**Figure 1.**
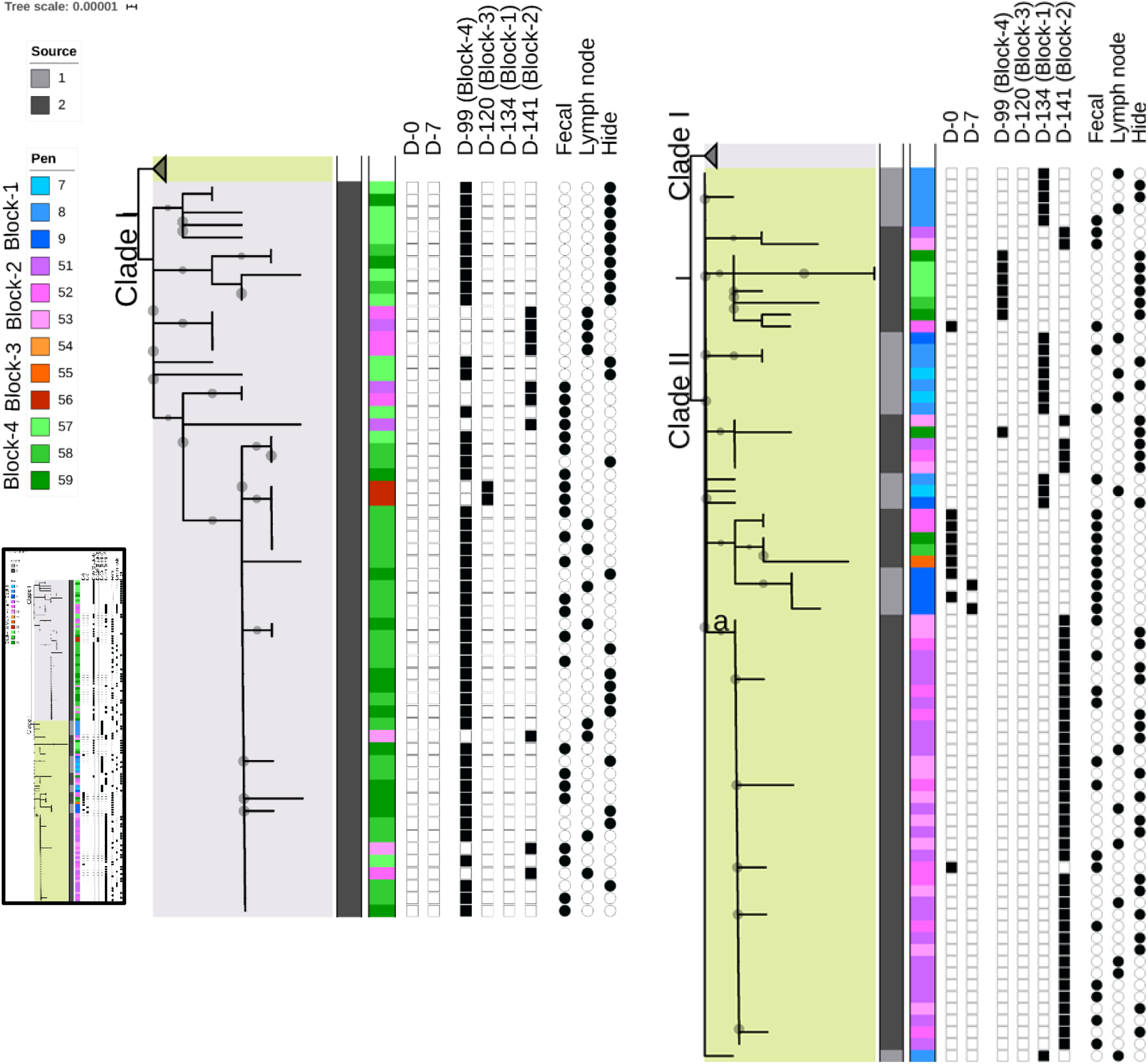
Phylogenetic analysis of 136 *Salmonella* Lubbock isolates, based on maximum-likelihood analysis of 84 SNP sites; of these, 36 were identified as parsimony informative and the remaining 48 were singleton sites. The tree scale (0.0001) was calculated to be equivalent to approximately 2.8 nucleotide substitutions per site. Full tree is presented in a miniaturized boxed view; expanded here, Clade I is in the middle and Clade II is on the right. Those branches with bootstrap support values of 80-100% are represented with a grey circle located in the middle of the corresponding branch and sized proportional to the given support values. Pen (first column) and cattle source (second column) colors are presented in the legends. Sampling days are presented in the next six columns. Sample types are indicated in the last three columns.

The cgMLST allelic distance measured among isolates of Lubbock (n=136) found in this study was 10 or less. The most prevalent clonal group was identified in the HC0_214521 cluster (n=31) and this clonal group was only observed at slaughter, and only among pens that were from Source 2; further, 26 of the isolates in this group were from Block 4. Another clonal group of the HC0_214545 cluster (n=28) was observed only in Block 2 and only at slaughter. Interestingly, isolates in Cluster HC0_215282 (n=9) were only observed in cattle in pens from Block 1 at slaughter and identified once again from Block 2 at slaughter. Isolates that were identified in Cluster HC0_214569 (n=2) were only observed in Pen 9 during the early feeding period (Days 0 and 7) and were not identified later during the study period (Fig. S1a). There were no unique patterns observed among different sample types (i.e., fecal, lymph node, and hide). The important features contributing most to the clonality of Lubbock were ordered from highest to lowest as experimental block, pen, source, day, and lastly sample type based on their ReliefF scores (Table S2) which likewise confirmed the phylogenetic findings.

As of August 21, 2020, Lubbock isolated from this study were most closely associated with two Texas origin dairy cattle isolates found in EnteroBase at HC2 and HC5 levels, respectively. An additional two Texas origin isolates (beef cattle and beef product origin) were in the same HC10 cluster. At the HC20-level, there were 139 isolates which were from the USA – and mainly from the states of Texas and Kansas. Isolates in this cluster were derived mainly from cattle liver abscess and beef products; importantly, no human origin isolate was reported in this cluster.

### *S*. Anatum

The genome of a previously isolated Anatum strain (GenBank: CP007483.2) was selected as the reference genome; prophage regions (n=5, Table S1) were masked from this reference genome. One isolate in this study (BioSample ID SAMN10910080) showed high divergence (> 5000 SNP differences as compared to the reference genome) was unable to be genotyped using the McCortex genotyping algorithm; therefore, this isolate was excluded from further analysis. As a result, 112 isolates were carried forward for SNP analysis. The best-fit nucleotide substitution model was selected as K3P+ASC according to the lowest BIC value. The resulting tree contained two distinct clades (Clade I and II)containing well-supported subclades (bootstrap values of >80%) that showed distinct patterns relating to source, day, experimental block, and pen. While isolates derived at slaughter from cattle in Pens 7, 8, and 9 (Source 1, Block 1) were observed only in Clade I, isolates from cattle in Pens 55, 56, 57, and 58 (Source 2) were observed only in Clade II. When sample day-related phylogenetic relationships were explored, no isolates were observed in Clade I from Day 7 and no other sample day-related patterns were apparent for either clade (Fig. 2).

**Figure 2.**
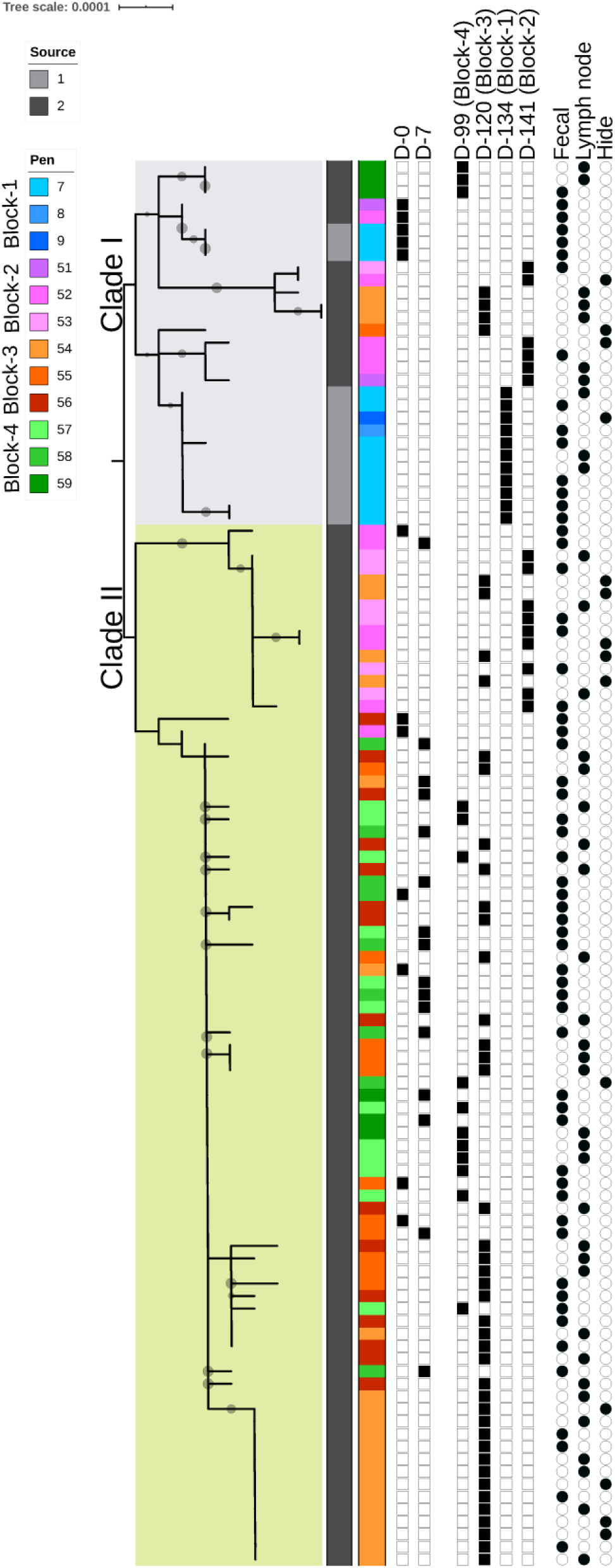
Phylogenetic analysis of 112 *Salmonella* Anatum isolates based on maximum-likelihood analysis of 66 SNP sites; of these, 36 were identified as parsimony informative and the remaining 30 were singleton sites. The tree scale (0.0001) shows approximately 2.2 nucleotide substitutions per site. Those branches with bootstrap support values of 80-100% are presented with a grey circle located in the middle of the corresponding branch and sized proportional to the given support values. Pen (first column) and cattle source (second column) colors are presented in the legends. Sampling days are presented in the next six columns. Sample types are indicated in the last three columns.

Considered in their entirety, Anatum isolates from this study did not show a cgMLST allelic distance of more than ten, except one isolate (same isolate that was excluded from the SNP analysis) that showed a large cgMLST allelic distance (50 or less) compared to the rest of the isolates at HC50 cluster (Fig. S1b). The most prevalent clonal group of Anatum was HC0_214518 (n=34), which was only observed in Blocks 3 and 4 and from Day 0 until the end of the study (Day 141). The second most prevalent (n=13) clonal group was HC0_214565, which was only observed in Pen 54 at slaughter. The clonal group of HC0_215273 (n=11) was only observed in Pen 52 during the feeding period on Day 0 (n=1) and Day 7 (n=1); at slaughter, this clonal group had become prevalent in the rest of the pens from Source 2 (n=9). This cluster was never identified in cattle from Source 1 during the study period. In contrast, the HC0_215308 clonal group (n=8) was only identified at slaughter and only in cattle from Source 1 (Fig. S1b). Based on the ReliefF scores, experimental block was the most important feature for explaining Anatum clonal differences, followed by pen, source, day, and sample type (Table S2).

The Anatum population identified in this study did not cluster with any public *Salmonella* genomes until the HC50-level. As of August 21, 2020, there were 209 *Salmonella* genomes that shared the same clustering at HC50. These genomes mainly originated from Texas and Kansas, USA and were derived from cattle and beef product samples. Of interest, the single Anatum isolate that was highly divergent when compared to the rest of the Anatum population was found to be in the same HC2 cluster as a single equine isolate derived from Texas in 2015.

### *S*. Montevideo

A previously sequenced *S.* Montevideo (GenBank: CP032816.1) strain was the genetically closest genome to the Montevideo isolates derived from our study. Prophage regions (n=4) masked in the reference genome are presented in Table S1. The tree was inferred with a K2P+ASC model (according to the lowest BIC value) and resulted in two clades (Clade I and II) for 68 Montevideo isolates. Clade I contained only a single distinct hide origin isolate derived from a steer located in Pen 52. The remaining isolates all were in Clade II. There was a subclade (a) within Clade II (Fig. 3) that harbored only isolates derived at slaughter age though from all sample types. The majority of isolates in this subclade was from Source 1, especially from Pens 8 and 9. The remainder of the subclade was consistently populated with isolates derived from feces sampled during the early feeding period, from a variety of pens. Montevideo phylogeny revealed a highly conservative tree with distinct day-(or sample type), and pen-related patterns and resulted in the lowest number of SNPs (n=12) when compared to the other serotypes.

**Figure 3.**
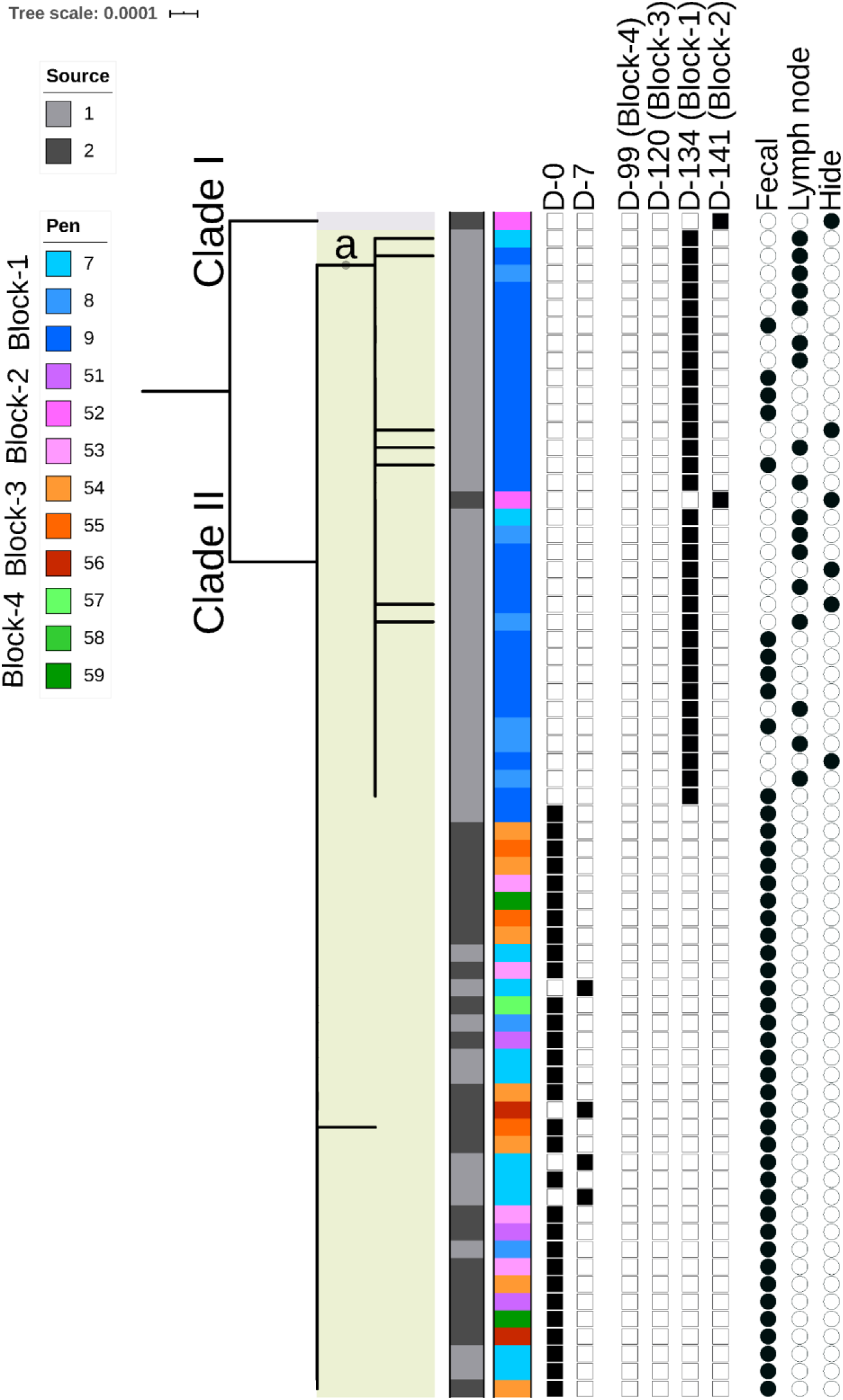
Phylogenetic analysis of 68 *Salmonella* Montevideo isolates based on maximum-likelihood analysis of 12 SNP sites; of these, one was identified as parsimony informative and the remaining 11 were singleton sites. The tree scale of 0.0001 is equivalent to approx. 0.4 nucleotide substitutions per site. Those branches with bootstrap support values of 80-100% are presented with a grey circle located in the middle of the corresponding branch and sized proportional to the given support values. Pen (first column) and cattle source (second column) colors are presented in the legends. Sampling days are presented in the next six columns. Sample types are indicated in the last three columns.

Montevideo isolates (n=68) identified in this study were highly clonal and isolates were no more than two cgMLST allelic distance values removed from each other. The most prevalent HierCC clonal group was HC0_214530 (n=30), which was observed only in the early feeding period (Day 0 and 7) in all pens (except Pen 52) and was derived from fecal samples of cattle that are from both sources. The second most prevalent clonal group was HC0_215274 (n=22), which was only observed among pens from Block 1 (Source 1) at slaughter, though it was found across all types of samples (Fig. S1c). Sampling day was the most important feature with the highest ReliefF score for clonal clustering of Montevideo, followed by sample type, pen, source, and experimental block (Table S2).

Montevideo isolates from this study were found in the HC20_9091 cluster along with a single isolate derived from swine feces in Texas. Besides that, isolate, there were no other similar isolates observed in the same cluster until the HC50-level. At HC50, a total of 1,543 isolates was found to cluster with our isolates, and the majority of those isolates originated from the USA (mainly, the states of Texas and California). Isolates in this cluster were primarily derived from beef products, followed by other cattle sources, including the feedlot environment and human clinical samples.

### *S*. Cerro

The Cerro population identified in this study was never observed in fecal samples from the early feeding period; thus, the analysis was restricted to slaughter age samples. The complete genome of a Cerro strain (GenBank: CP008925.1) was selected as the reference genome, and five prophage regions (Table S1) were masked before SNP analysis. The phylogenetic tree of 64 Cerro isolates was inferred using TIMe+ASC nucleotide substitution model according to the lowest BIC value. The resulting phylogenetic tree revealed two separate clades (Clade I and II). Among those, Clade I was well-supported (bootstrap value of > 80%) presenting distinct source, slaughter sampling day (collinear with experimental block), and pen patterns (Fig. 4). Isolates derived from cattle in Pens 7, 8 and 9 from Block 1 (Source 1 and slaughter Day 134) and Block 4 (Source 2, slaughter Day 99) were found only in Clade I. Clade II harbored only the isolates from Source 2 and mainly were from Pens 55 and 56 of Block 3 (slaughter Day 120). There were no obvious sample type-related phylogenetic patterns observed among Cerro isolates.

**Figure 4.**
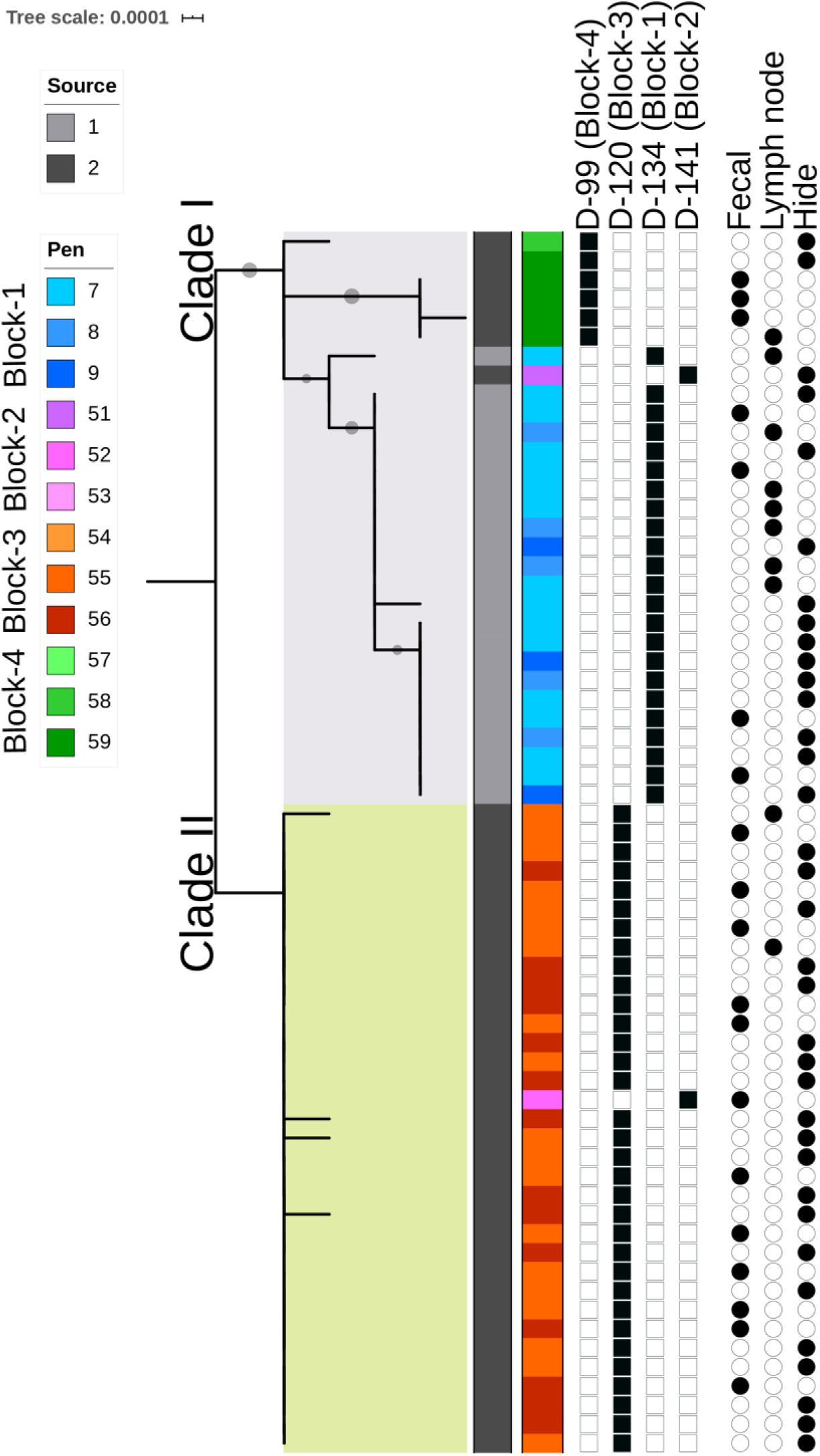
Phylogenetic analysis of 64 *Salmonella* Cerro isolates based on maximum-likelihood analysis of 17 SNP sites; of these, 9 were identified as parsimony informative and the remaining 8 were singleton sites. The tree scale (0.0001) shows approximately 0.4 nucleotide substitutions per site. S. Cerro isolates were derived only at slaughter age in this study. Those branches with bootstrap support values of 80-100% are presented with a grey circle located in the middle of the corresponding branch and sized proportional to the given support values. Pen (first column) and cattle source (second column) colors are presented in the legends. Sampling days are presented in the next four columns. Sample types are indicated in the last three columns

A total of three clonal HC0 clusters was identified with an cgMLST allelic distance of five or less. The most prevalent clonal HierCC cluster was HC0_214520 (n=23) which was only found in cattle from Source 2, and 22 of these were arose from Pens 55 and 56 (Block 3); the remaining isolate was derived from Pen 52. The second most prevalent clonal group was HC0_215280 consisting of 22 isolates; of those, 19 were derived from Block 1 (Source 1) while the remaining three isolates were from pens housing cattle from Source 2 (Fig. S1d). There were three isolates in the third clonal group of HC0_214566; all arose from samples taken from Pen 59 at slaughter. This group (HC0_214566) of isolates also differed from the rest of the isolates at the HC5 level (i.e., no more than cgMLST allelic distance of five). Based on ReliefF scores, the most important feature of clonal Cerro clustering was experimental block (collinear with slaughter day) followed by source, pen, and sample type (Table S2).

Similar to Anatum isolates, Cerro isolates were not found to be in the same cluster as any other public *Salmonella* genomes until the level of HC50. At the HC50 cluster, as of August 21, 2020 there were 283 matching genomes (HC50_996) and the majority of these were sequences from isolates collected in the United States and a few other countries (e.g., Taiwan and the United Kingdom). The isolates from the USA mainly were from Texas, followed by California. Cerro genomes at the HC50 cluster level were mostly associated with beef products, cattle, and humans.

### Other serotypes

Kentucky (n=11) isolates were derived only from early feeding period (Days 0 and 7) fecal samples of cattle purchased from Source 1 and were highly clonal; of these, 9 isolates were identified as being in a single highly clonal group (HC0_214531) as shown in Fig. S1e. Kentucky isolates were not closely related to any other isolated Kentucky genomes until the HC50-level. At this level (HC50_16), a total of 4,521 Kentucky genomes were matched with the HierCC inquiry, and mainly originated from the USA; of those, the majority of isolates were derived from California, followed by Georgia and then Texas. A majority of Kentucky genomes at the HC50 clustering level were associated with poultry, and rarely with either cattle or human isolates.

Newport (n=6) isolates were derived only from fecal samples and lymph nodes of cattle located in Pen 53 and only at slaughter age. These isolates were indistinguishable from each other and belonged to a single clonal group (HC0_215270) as also shown in Fig. S1f. Newport isolates derived in our study were no more than cgMLST allelic distance of 20 (clustered at HC20) from two *Salmonella* genomes derived from human clinical specimens in the USA. The next HierCC cluster match was observed at HC50-level with 284 isolates, the majority of those were from Arizona, followed by California and Texas. Isolates from this cluster were primarily isolated from human, seeds and vegetables and rarely found in cattle related sources.

Only a single Norwich isolate was identified at slaughter from a fecal sample of a steer located in Pen 52. This isolate was no more than five cgMLST allelic distance to four human isolates from the USA. In addition, there was a total of 72 human or outbreak origin Norwich genomes that were observed in the same HC10 cluster. There was no other public *Salmonella* genome found in the HC20 or HC50 cluster of the single Norwich isolate.

### Antibiotic resistance, plasmids, and pathogenicity island profiles

Among 399 *Salmonella* isolates screened for antibiotic resistance genes and plasmids. All Montevideo isolates (except for one) harbored a tetracycline resistance gene *tet*(A), all of which were located on an IncN (IncN_1) plasmid-related contig. A subset of isolates (n=62) carried an IncI1 (IncI_1_Alpha) plasmid, which was not serotype-specific and was found only among a subset of Anatum, Lubbock and Montevideo isolates.

A total of ten *Salmonella* pathogenicity islands (SPIs) were identified across these same 399 isolates. Of those, SPI1, SPI2, SPI5, and SPI9 were found in all serotypes with few exceptions; on the other hand, C63PI, SPI3, SPI8, SPI13, and SPI14 were found to be highly serotype-specific. The SPI4 mainly varied across the Cerro and Lubbock subpopulation of isolates, regardless of the clonal distribution of these isolates and of any other features such as pen, day, or sample type. Antibiotic resistance genes, plasmids and SPIs that were identified within each serotype are provided in Table 3. No distinct plasmidal or pathogenicity island profiles were found to be related to a specific clonal group (Supp. Dataset 1).

**Table 3.**
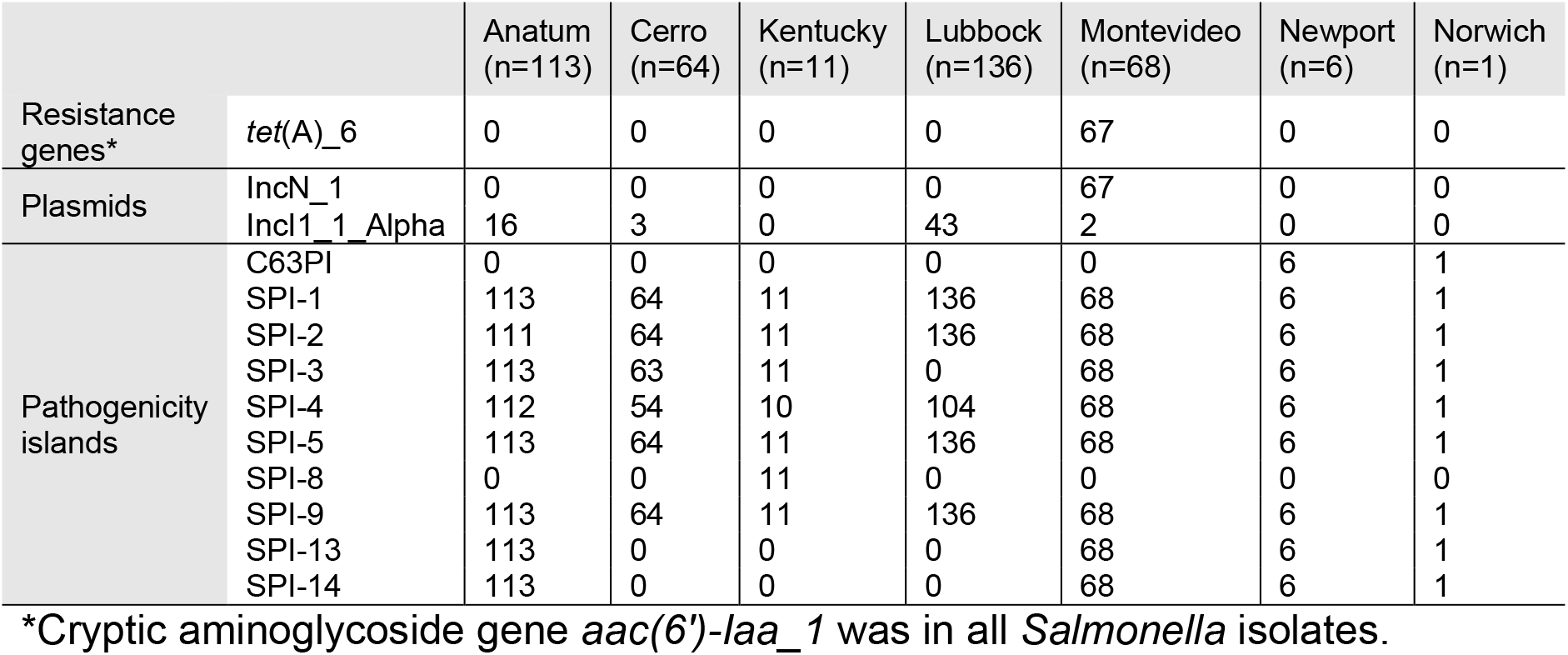
Distribution of isolates harboring antibiotic resistance genes, plasmids, and pathogenicity islands by serotype. Resistance genes were identified using the ResFinder Database updated on 06-03-2020 with the minimum threshold set for 97% for ID and 97% for coverage match. Plasmids were identified through the PlasmidFinder database updated on 06-03-2020 with the minimum threshold set for 97% for ID and 80% for coverage match. Pathogenicity islands were identified through the SPIFinder database updated on 07-24-2017 with the minimum threshold set for 97% for ID% and 60% for coverage match.

## Discussion

Our previously published randomized controlled longitudinal feedlot trial was initially designed to explore the effects of single-dose antibiotic metaphylaxis for bovine respiratory disease on antibiotic resistance and the prevalence of *Salmonella* in cattle feces during the feeding period, and in feces, sub-iliac lymph nodes and on hide surfaces at slaughter. The results of our previous study reflected no significant differences between the prevalence of *Salmonella* in cattle that received either antibiotic treatment (ceftiofur or tulathromycin) and those in the control group that received neither. What varied instead, was the distinct serotype distribution patterns observed across sampling days and sample types among the 12 pens, four experimental blocks, and the two sources the cattle were purchased from, which merited further investigation into how the subpopulations within the serotypes were influenced when a higher resolution was obtained. The isolates identified as Anatum, Lubbock, Cerro, Montevideo, Kentucky, and Newport serotypes each appeared in their own same legacy 7-gene MLST groups, which initially suggested clonal dissemination of *Salmonella* within the cohorts of cattle throughout the feeding period and at slaughter (25). However, initial examinations of the phylogenies of these longitudinally sampled isolates suggested that simplistic reasoning was likely wrong.

In this follow-up study, we further explored the clonal distribution and evidence for temporal and geographical dissemination of *Salmonella* isolates by performing core-genomic comparisons with increased resolution using core-genome SNP-based maximum likelihood phylogeny and allele-based approach using HierCC method. HierCC analyses directly supported the clonal distributions that were observed on the phylogenetic trees. The latter analysis method was used specifically to identify clonal groups, predict the importance of the study design-related features as well as the genomic distance between the isolates derived from this study and the isolates derived from clinical human cases. HierCC methods have been shown to be highly reliable in predicting legacy 7-gene MLST groups (HC900), endemic persistence of strains (HC100 and HC200), as well as the clonal groups of epidemic outbreak strain (HC2, HC5, and HC10) clusters based on distances observed among cgMLST alleles (38).

Overall, the results we obtained from SNP- and allele-based distance analyses showed that experimental blocks – wherein 11-12 cattle randomized to three adjacent pens were geographically co-located – and the geographical origin of the cattle (source) were the most consistently predictive factors explaining clonal clustering and expansion observed in the *Salmonella* population. These two factors were followed by sampling day and sample type, though of lesser importance.

*Salmonella* Cerro (legacy 7-gene MLST ST367) isolates were identified in cattle from all experimental blocks; however, this serotype was only isolated from samples collected when cattle were eligible to be sent to slaughter between Days 99 and 141 (specific slaughter sampling dates by experimental block are listed in Table 1). Our findings show that clonal clustering/expansion of the Cerro population was most influenced by experimental block (or, by slaughter day due to collinearity), followed by source, pen, and sample type. However, it seems most likely that the clonal effects observed are attributable to block rather than sampling day. Relatively short intervals between sampling days at slaughter eligibility underlies this hypothesis, which will also have applied to the other serotypes that were derived during the study within the constrained period of slaughter, or where the clonal expansion of *Salmonella* seemed affected less by day than by block. While the Cerro population derived from our study were genetically clustered with other publicly available genomes of the isolates originated from Texas, the beef-, human-, and cattle-origin Cerro population showed a strong endemic persistence of the outbreak-related strains and had no more than 50 cgMLST allelic distance from the isolates derived from our study. Preceding our study, the work of Kovac et al. (2017) identified geographical location as a potential driver of the clonal expansion of the Cerro population (ST367) derived from dairy cattle located either in Texas or New York state (52).

Prior to this present study, the potential drivers of the clonal clustering/expansion of Lubbock and Anatum populations in cattle had not been explored. Our results indicate that experimental block, pen, and cattle source are the most important features for explaining clonal relatedness of Lubbock (ST413) and Anatum (ST64) populations in fed cattle during a study period such as ours. In addition, the HierCC results of our study suggest that dissemination of Lubbock and Anatum related clones has been restricted primarily to Texas and Kansas, U.S. states where these two serotypes mainly were isolated from cattle and beef products. However, neither set of serotypes clonal groups have been associated with human isolates at the HC50-level. An emerging serotype first reported in 2015, Lubbock was first identified in a sub-iliac lymph node of a harvest-ready steer located in Texas. The study of Bugarel et al. (2015) suggested the novel serotype evolved from its ancestral serotype Mbandaka through recombination events (75). As of yet, no Lubbock strain was reported as a source of human clinical *Salmonella* outbreaks. Thus, the possible public health outcome of the Lubbock population remains unknown.

In contrast to *S*. Anatum, Lubbock and Cerro, the clonal clustering/expansion of the Montevideo (legacy 7-gene MLST ST138) population was influenced mainly by sampling day, sample type and pen; in contrast to the other serotypes, experimental block, and cattle had much less influence on their apparent dissemination. This difference was perhaps not surprising since the majority of Montevideo strains were identified in fecal samples initially from the beginning of the early feeding period and across all pens from both sources or mainly from Source-1 at slaughter time on day 134 (Fig. 3). While our machine-learning approach suggests that features related to sampling day and sample type had more bearing on Montevideo clonal clustering/expansion, it also is possible that our study design might have masked the pen-, block-, and source-related influence. The well-supported (bootstrap value of >80%) subclade (a) within the Clade I observed in the phylogenetic tree of Montevideo also reinforced that there was an even greater effect of pen when multiple sample types were accounted for (Fig. 3). In addition, sample type has not earlier been reported as being associated with clonal clustering/expansion of *Salmonella* subpopulations reported (42, 51). Typhimurium and Derby isolates derived from swine feces, lymph nodes and carcasses in farms located in the state of North Carolina have yielded, for example, no phylogenetic relations when comparing among the sample types from which isolates were derived (42). The Montevideo (legacy 7-gene MLST ST138) population in our study displayed more similarities with previously recorded cattle-origin isolates than with human-origin isolates at the HC50-level. The research of Nguyen et al. (2018), likewise, revealed with a few exceptions that Montevideo legacy 7-gene MLST ST138 strains were related mainly to cattle origin isolates and were located in different clades than human outbreak origin isolates which mainly were from ST316 (43).

We also identified 11 highly clonal Kentucky isolates from the early feeding period (Day 0 and 7); importantly, these were identified only from those cattle purchased from Source 1. We believe that Kentucky was the serotype most likely to have pre-existed in a group of cattle prior to placement in the feedlot. Further, this helps to explain the apparent greater effect of source and sampling day than of experimental block and pen (geographical and environmental surrogates) on the clonal clustering/expansion of the Kentucky population. In addition, a total of six Newport isolates were identified, all of which were found to be highly clonal and originated from either a lymph node (n=5) or fecal sample (n=1) of cattle housed in the same pen (Pen 53). This population exhibited no more than 20 cgMLST allelic distance from human-and outbreak-related isolates. Although these isolates had no antibiotic resistant genes, the presence of the highly clonal Newport population in the lymph nodes poses a potential risk for contamination of ground beef. This is a tangible public health risk, since a recent *Salmonella* outbreak similarly originated in a contaminated ground-beef product, was traced to a pan-susceptible Newport isolate in 2018, and resulted in 403 reported cases in 30 states causing 117 hospitalizations (76). Besides Newport, our study identified a single Norwich strain isolated from a cattle fecal sample at slaughter. This isolate was found to be related to 70 human and outbreak origin isolates at the HC10-level and 4 out of 70 came from the same cluster at the HC5-level which is reveals this strain to be a clonal epidemic outbreak strain (38).

Our data did not reflect any SPIs specific to a certain clonal group of *Salmonella*. The exception was the presence of SPI4, which varied within both the Cerro and Lubbock populations. The SPI4 is known to carry virulence genes responsible for epithelial adhesion and host invasion (77). When we examined the absence or presence of SPI4 regions at the population level, for instance, patterns related to individual cattle seemed more consistent compared to those related to pen, block, source, and day. Over the entire feeding period, we identified only two types of plasmids (IncN and IncI1) – both, incompatibility group plasmids previously reported to harbor beta-lactam class resistance genes in *Salmonella* isolated from farm animals (78). Even though the selection pressure of ceftriaxone metaphylaxis was applied to one group of cattle on day 0, none of the isolates derived bore IncI1 and IncN plasmids harboring *bla* genes. The presence of the IncI1_alpha plasmid was related neither to a particular clonal group nor to a particular serotype. IncI plasmids can, in fact, be found in a wide range of *Salmonella* serotypes and may or may not carry antibiotic resistance and virulence genes; however, the exact function of IncI1 plasmid remains unknown (79).

We found an IncN_1 plasmid and *tet*(A) gene associated with all of our Montevideo isolates. Cattle-origin Montevideo isolates were previously found to be related to an IncN_1 plasmid harboring *tet*(A) genes (28); however, no sequence type (legacy 7-gene MLST) of the Montevideo isolates in that study were reported. Another study – Nguyen et al. (2018) – examined a cattle origin Montevideo (legacy 7-gene MLST ST138) population, but did not report any IncN plasmids and tetracycline resistance genes (43), suggesting that the IncN plasmid and *tet*(A) gene presence in Montevideo ST138 may be related to the particular feedlot environment or the geographical origin of the cattle. We have previously described the antimicrobial susceptibility phenotype of these same isolates (25) and showed that the Montevideo isolates that carried the *tet*(A) gene were also found to be phenotypically resistant to tetracycline. However, based on sequencing data, 21 isolates that were previously identified as phenotypically resistant to streptomycin did not bear any such streptomycin (aminoglycoside) resistance genes, such as of the *str*, *aad*, *aph,* or *arm* families. This inconsistency between the genotypic and phenotypic streptomycin resistance could possibly be ascribed to the uncertainty of the MIC breakpoints that have been determined for phenotypic streptomycin resistance (80).

Our results showed no antibiotic resistance, plasmid, and pathogenicity island profile association among varied *Salmonella* subpopulations derived from individual cattle, pens, blocks, and sources during the study period. However, the prophage regions of the isolates that are expected to usually vary by host and environment was not explored in our study. Beside the lack of prophage profile findings, another lack of finding was the absence of samples from the environment (e.g., pen floor, feedlot dust) to measure the environmental contribution on *Salmonella* dynamics in feedlot cattle.

We employed a supervised machine-learning algorithm to estimate the relative importance of features such as source, experimental block, pen, sampling day, and sample type for explaining clonal clustering/expansion of *Salmonella* serotypes. We used a Random Forest algorithm with ReliefF-based feature selection to rank the prediction power of these features by their potential to forecast clonal outcomes (68, 71). While the outcomes of the feature importance were consistent with the core-genomic subpopulation-related analysis performed in this case, the interpretation of results in cases where machine-learning methods are used needs to be handled with care; specifically, with an eye to prioritizing the qualitative roles of biology and study design over numeric predictions. Our study suggests that the application of machine-learning tools, when used for the prediction of feature importance on clonal clustering and expansion, can be a promising approach. However, used in studies such as ours, with more limited sample size numbers, makes interpretation in relation to the biology and study design features even more important.

In summary, this was the first study of its kind to attempt to monitor and measure the microevolutionary progress of *Salmonella* subpopulations over time among various sample types collected from a group of beef cattle during the feeding period and at slaughter. We aimed to explore the potential initiators and drivers of *Salmonella* population dynamics as observed in feces, lymph nodes, and hide samples, all of which are among potential contaminants of beef products at slaughter. Our research findings stress the importance of cattle origin, which impacts background microbiota on arrival and also the starting feedlot environment into which these same cattle are placed (including the geospatial placement of cattle and their pens). In total, these two features are the main drivers of initial shared *Salmonella* clonal profiles that later morph in response to microevolutionary pressures. In order to combat antibiotic resistance and reduce the pathogenicity of *Salmonella* found in final beef products, more research is needed to explore the feedlot management and environmental effects on clonal *Salmonella* clustering and expansion, including pre-existing soil and manure pack microbiota and dust contribution as well as the bacteriophages that are in the feedlot environment.

## Acknowledgments

We acknowledge the H. M. Scott laboratory graduate and undergraduate students for assisting with sample collection and microbiological processing. This study was funded by the National Cattlemen’s Beef Association, a contractor to the Beef Checkoff (no. 22615).

